# TFound: A Software to Map Cis-Regulatory Elements in Yeast

**DOI:** 10.1101/474957

**Authors:** Adriano Gomes-Silva, Rafael Silva-Rocha

## Abstract

Transcription factors (TFs) are responsible for regulating the rate of transcription of genes in all organisms. These factors can be represented computationally by position weight matrices (PWM). TFound was developed to allow the detailed visualization of predicted binding sites of transcription factors in multiple sequences based on the PWMs, by using the graphic user interface (GUI). The tool was loaded with the genome of *Saccharomyces cerevisiae* and PWMs from the YeTFaSCo database (http://yetfasco.ccbr.utoronto.ca/), also allowing the insertion of new sequences and PWMs. Thus, the user is allowed to load custom PWMs and genomes to perform easily mining and visualization of binding site motifs of *cis*-regulatory elements of interest, permitting an efficient way to inspect DNA assembly projects for complex synthetic circuits. This work describes the functionality of the current version of the tool, which is coded in *Python* and is freely available at the repository https://github.com/adri4nogomes/TFound.

## MOTIVATION

The ability to coordinate the expression of thousands of genes by processing a wide range of environmental inputs is a fundamental feature of any living organism ^1^. To decipher the logic underlying such complex regulatory network is crucial to understand the basic signaling principles of life machinery, providing knowledge to reprogram it to execute novel synthetic functions ^2,3^. However, despite being essential to engineering, the inference of such complexity remains a major bottleneck, especially to eukaryotes ^4–7^. Moreover, Fungi are a powerful resource for biotechnological applications, from biomass conversion aiming biofuels production to the synthesis of industrially relevant compounds ^8,9^.

In particular, microorganisms of the yeast group are considered to be key providers of several biotechnological solutions. In recent years, numerous studies have focused on the engineering of synthetic circuits in yeasts aiming to expand the metabolic and biosynthetic capacities of these organisms and to increase their tolerance to aggressive environmental conditions such as those related to process-specific industrial parameters ^10,11^. Yet, when several biological parts are combined together, novel functional elements can be unintendedly generated ^12,13^. These new features could interfere with the function of the final circuit, leading to the disruption of the desired behavior. Therefore, it is imperative to check the final assembled circuits aiming to identify undesired regulatory elements generated during the assembly ^12,14^. In addition to this application, it would also be interesting to have a tool to easily identify conserved *cis*-elements in related species. Examples could be the identification of *cis*-elements for known TFs from *S. cerevisiae* in other yeast or filamentous fungi, which could lead to the finding of orthologous TFs genes in the organism of interest ^4,15,16^.

In this context, here we present a user-friendly and integrated platform to infer *cis*-regulatory elements and its patterns of distribution in a genome-scale approach, from data selection to visualization. The program is initially loaded with the genome and annotation of *S. cerevisiae* S288C and with the set of position matrices of The Yeast Transcription Factor Specificity Compendium (YeTFaSCo) ^17^, but it supports the receipt of other annotated genomes and sets of position matrices. We hope this new tool could be useful for the Synthetic Biology community, aiming the inspection of *in silico* designed synthetic circuits to avoid the generation of unwanted regulatory elements.

## SEQUENCE INPUT

TFound was developed in *Python* and its graphical interface developed implementing Tkinter and Matplotlib libraries. The system allows as input DNA sequences from the *S. cerevisiae* genome (which is already preloaded in the current version of the program), from FASTA files or sequences typed directly into the given field. Also, it accepts the entry of other genomes using FASTA files plus a GFF file for the annotation (**Fig. 1A**). Once a new sequence in FASTA is inserted, a list of sequences is presented in a table next to their respective IDs. In genome insertions or in the use of the pre-loaded genome of *S. cerevisiae*, the sequences are listed next to their respective gene names (**Fig. 1D**). Genes can be filtered by their descriptions (**Fig. 1B**) and, in the case of the preloaded genomes, they can also be filtered by regulators from which they are targeted (**Fig. 1C**). When using entire genomes, it is possible to choose the quantity of upstream and downstream bases to be analyzed (for a typical analysis, we select 1000 bp upstream and 100 bp downstream from the gene ATG, **Fig. 1F**). Finally, filtered sequences and their specified upstream and downstream regions can be exported in the FASTA format (**Fig. 1E**).

**Figure 1.**
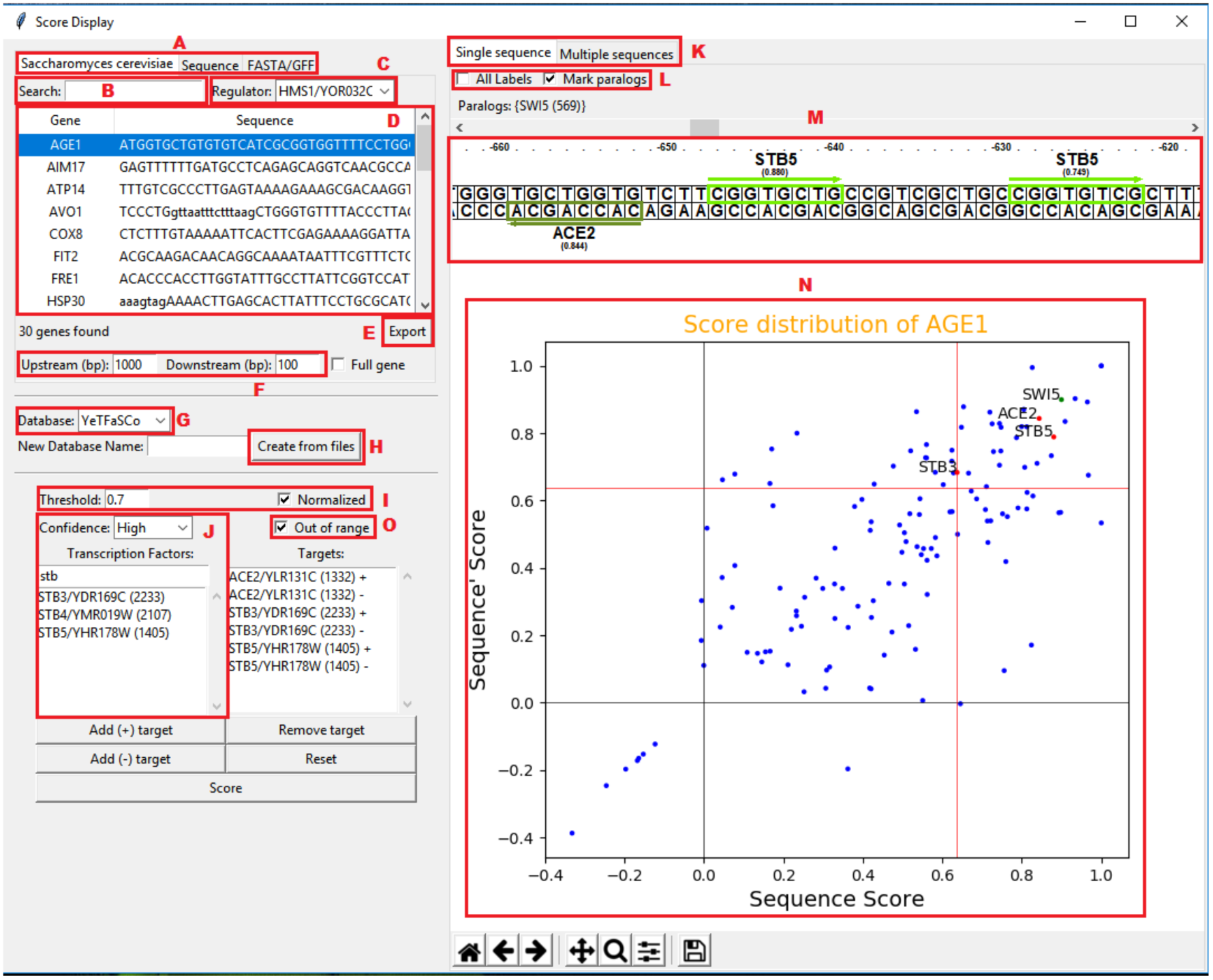
Program interface searching the ACE2, STB3 and STB5 transcription factors in single sequence mode for the *S. cerevisiae* AGE1 gene. (A) Input options. Search by (B) Gene/ID and (C) regulator. (D) Table of sequences. (E) Export sequence’s slice to FASTA. (F) Gene slice selector. (G) Database selection. (H) Database creation. (I, O) Transcription factors options. (J) Transcription factors search. (K) Visualization selection. (L) Single sequence options. (M) Sequence detailed view. (N) Score distribution of sequence.

## TRANSCRIPTION FACTOR INPUT

Position weight matrices (PWMs) are used to represent binding sites motifs of transcription factors. Initially, loaded PWMs from the YeTFaSCo collection (**Fig. 1G**) are listed for filtering and selection, but new PWMs can be inserted by creating a new database (**Fig. 1H**). The user can filter the PWMs for selection by trust level, default name, systematic name, or ID (**Fig. 1J**). By selecting a PWM, it is possible to add it for analysis in the sense and/or antisense strand. It is also necessary to determine the score threshold for the labeling of the transcription factors in the sequence, as well as to determine whether the score will be normalized or not (**Fig. 1I**). Finally, the user choses whether bases not determined at the end of the sequences should be considered in the tags (**Fig.1O**).

## OUTPUTS

TFound has two output types: single sequence and multiple sequences (**Fig. 1K**). Using a single sequence, the output is the detailed sequence bases with the labeling of the positions where the selected transcription factors exceed the selected threshold score. This is presented in the sense and antisense strands together with the respective TF names and scores, which allows inferring the binding sites, correlations and conflicts between TFs (**Fig. 1M**). A graphic distribution of the maximum scores (at sense and antisense strands) is also displayed for each TF along with the selected confidence level (**Fig. 1N**) and the user can interactively label only selected or all TFs in the database (**Fig. 1L**). This allows the visualization of TFs with higher score in the sequence and its direction, and also the identification of paralog ones (**Fig. 1L**).

In the selection of multiple sequences (**Fig. 2**), the positions of the selected TFs (**Fig. 2B**) exceeding the threshold score for each sequence is presented, allowing the visualization of the distribution of TFs at the positions of the sequences. It is also possible to filter only for sequences in which some (or all) of the selected TFs appear (**Fig. 2A**), allowing the visualization of cooccurrence of binding sites in the sense (**Fig. 2C**) or antisense (**Fig. 2D**) strands. Finally, the list of target genes is presented in the graphical form (**Fig. 2E**) and the position relative to the ATG (**Fig. 2F**) is presented.

**Figure 2.**
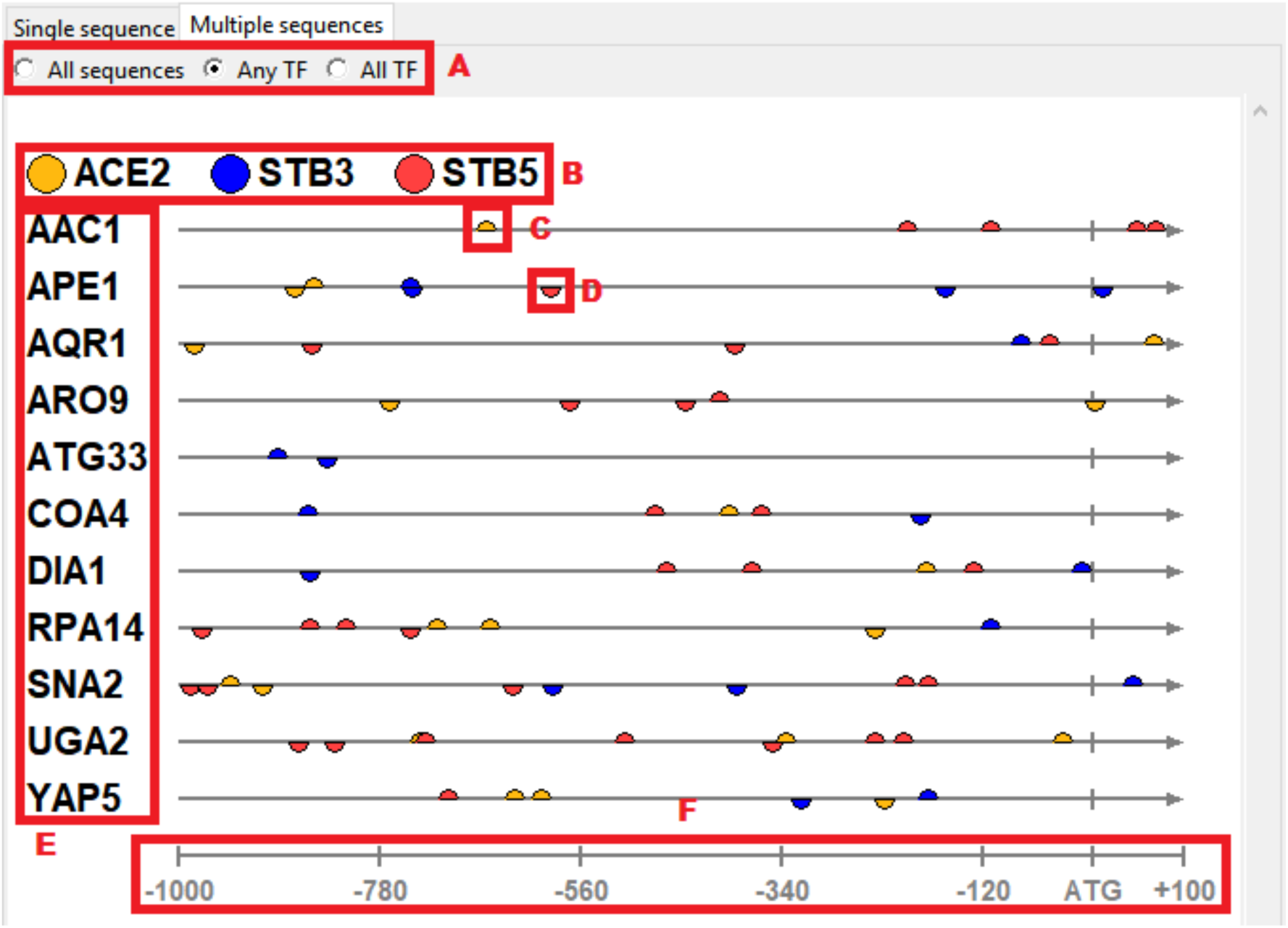
Search by transcription factors ACE2, STB3, and STB5 in the sense and antisense tapes in the upstream and downstream regions of multiple sequences, displaying only the sequences that show the presence of some of the selected transcription factors. (A) Transcription factors options. (B) Transcription factors selected. (C) Sense and (D) antisense binding sites. (E) Gene targets. (F) Sequence ruler.

## CONCLUSIONS

From any genome and its annotation, TFound provides an easy selection of regulatory regions or a set of FASTA sequences for further analysis. Based on the Position Matrices of transcription factors (e.g., Position Weight Matrix – PWM and Position Frequency Matrix - PFM) ^7^, TFound offers a robust analysis of scores distribution in positions at sense and antisense strands in a whole genome scale or in selected sequences. From these analyzes, TFound display the patterns of distribution of TFs motifs, the correlation between sets of TFs, the correlation between TFs and their positions of greater probability of connection. Also, TFound provides a simple visualization of possible co-locations between TFs, generating graphs and tables that facilitate the display and understanding of such information. As for the initial version of the software, the user can investigate the *S. cerevisiae* regulatory machinery, using loaded reference genome, annotation and a set of position matrices from the YeTFaSCo collection. TFound is a modular platform and supports other annotated genomes and sets of Position Matrices, providing a simple and accessible tool to decipher multiple regulatory mechanisms through the inference of TF-*cis* regulatory element interaction, paving the way to the understanding and precise engineering of a range of microorganisms. In the future, we plan to change the graphical interface library for the GTK+ library to make it more user-friendly and functional, adding score and position distribution charts and including the search for binding sites using the IUPAC code, palindromic sequences, and other types of position matrices.

## AVAILABILITY

The project is open-source and has been completely developed in Python using the following libraries: BeautifulSoup for downloading the database; Pandas, Numpy, and BioPython ^18^ for data manipulation; Tkinter and MatPlotLib ^19^ for the development of the Graphical User Interface (GUI); Numba ^20^ for parallel processing of data. All scripts are available in the following https://github.com/adri4nogomes/TFoundrepository.

## ACKNOWLEDGMENTS

The authors are thanks to laboratory members for insightful discussion on the manuscript. RSR was supported by FAPESP Young Research Award (grant# 2012/22921-8). AGS was supported by a FAPESP Master Fellowship (grant# 2018/06509-6).

